# Computational psychiatry 2.0 and implications for stress research

**DOI:** 10.1101/483164

**Authors:** Anton A. Pashkov, Mikhail A. Berebin

**Author notes:** **Corresponding author:** Anton A. Pashkov, Biomedical school, Department of Clinical Psychology, South Ural State University, Chelyabinsk, Russia.

## Abstract

Stress-related disorders are highly prevalent in modern society and pose significant challenge to human’s health. Being recently emerged branch of psychiatry, computational psychiatry is geared toward mathematical modeling of psychiatric disorders. Harnessing power of computer sciences and statistics may bridge the complex nature of psychiatric illnesses with hidden brain computational mechanisms. Stress represents an adaptive response to environmental threats but, while getting chronic, it leads to progressive deflection from homeostasis or result in buildup of allostatic load, providing researches with unique opportunity to track patterns of deviations from adaptive responding toward full-blown disease development. Computational psychiatry toolkit enables us to quantitatively assess the extent of such deviations, to explicitly test competing hypotheses which compare the models with real data for goodness-of-fit and, finally, to tethering these computational operations to structural or functional brain alterations as may be revealed by non-invasive neuroimaging and stimulation techniques.

It is worth noting that brain does not directly face environmental demands imposed on human or animal, but rather through detecting signals and acting out via bodily systems. Therefore, it is of critical importance to take homeostatic and allostatic mechanisms into account when considering sophisticated interactions between brain and body and how their partnership may result in establishment of stress-susceptible or resilient profiles.

In this article, with a particulate focus on brain-gut interactions, we outline several possible directions to widen the scope of application of computational approach in mental health care field trying to integrate computational psychiatry, psychosomatics and nutritional medicine

## Introduction

Investigation of stress issue in modern science is one of the priority areas of research, primarily due to its high social significance. Chronic stress is a risk factor for a number of mental disorders, such as anxiety disorders, depression, PTSD, adjustment and eating disorders (1). In this regard, fruitful way of dealing with the issue draws on development of sensitive tools for stress vulnerability elicitation, investigation and development of novel treatment options.

Stress-related psychiatric disorders are characterized by impairments not only in the brain circuits, but also in brain-body interaction pathways, which are reflected in the very definition of stress as the organism’s response to an imaginary or real threat to physical or mental integrity (2). Considering a huge number of factors influencing development of stress-related disorders and the success of therapeutic interventions impose special requirements on the investigation of such processes in the body. Despite the notable progress in identifying genetic, molecular, cellular, network mechanisms of development of such disorders, it is much more difficult to combine the results of individual experiments into a single theory or set of theories (3). This integration has become possible with the introduction of computational models into psychiatric methodology (4). Thus, one of the promising ways to meet the challenge is the recent emergence of a new field of computational psychiatry, which has already proven its usefulness in search for new mechanisms of the pathogenesis of mental disorders.

### Stress research meets computational psychiatry

Computational psychiatry allows us to link together different levels of the description of studied phenomenon, identify key sub-parts or previously unknown constituents of cognitive functions and, in addition, relying on hypotheses being put forward, to create and test models, quantifying their parameters (for example, learning rate, confidence, prediction error) (5,6). Computational models enable one to account for a large scope of observations with relatively few parameters. Such mathematical implementations offer essential advantages, allowing for modeling non-linear interactions between different brain regions or between brain and body.

One should also keep in mind that nowadays psychiatry experiences a shift in the researchers’ focus of attention from categorical diagnostics to exploration of mental disorders’ spectra (autism, schizophrenia, obsessive-compulsive disorder). This new ground is being incorporated into the DSM-5 and upcoming ICD-11 (7). Conventional approaches to dealing with psychiatric disorders rest on descriptive diagnostic systems such as ICD10 or DSM-5, which are currently lacking of mechanistic power to explain symptoms in terms of genetic, epigenetic, molecular abnormalities and impairments in meso- and macroscale patterns of structural or functional brain connectivity (8). However, research domain criteria framework (RDoC) has now appeared to take advantages of state-of-the-art findings in biological psychiatry as well as to fix some flaws in disease classification guidelines being currently in use. The search for biomarkers is a leading endeavor in biological psychiatry and this line of action can be successfully embedded into the global framework of computational psychiatry (9).

A distinct branch of computational psychiatry consists of application of new methods of machine learning for solving problems of classification and regression in clinical practice. Joining these analysis techniques together with a plethora of data from international research collaborations and depositories has the potential to foster the biologically centered re-conceptualization of mental disorders. However, due to page restrictions, this article does not consider this research direction, but given subject has thoroughly been discussed elsewhere (10, 11).

Perhaps the most perspective framework to handle the immense complexity of brain inputs and outputs is predictive coding theory, the main idea of which is that our brain is constantly generating and updating hypotheses that predict sensory inputs at varying levels of abstraction (12).

Converging evidence within the field of computational neuroscience states that animals and humans are capable of extracting statistical regularities from environmental events. By tackling the probabilities, our brain is capable of making predictions of future outcomes (13). In what follows, these predictions should be compared with actual sensory input in different levels of neural hierarchies. In the case of discrepancy between them, prediction error is emerged and transmitted up the neural hierarchies. Highly relevant in context of stress research is the fact that uncertainty or volatility of environment poses a substantial challenge for mastering these predictions properly (14). It has been recently proposed that beliefs about uncertainty can mediate the strength of stress response (15).

According to findings derived from computational neuroscience experiments, various neuromodulators can differently affect uncertainty computations. All of these neuromodulators are being altered during stress conditions (16). Two key neurotransmitters which are able to influence such computations are norepinephrine (NE) and acetylcholine (Ach). Ach signals so-called expected uncertainty, i.e. unreliability of predictions based on environmental cues. Whereas NE is concerned with unexpected uncertainty, i.e. global alarm system which regulates context switching and novelty detection (17). The more NE and Ach levels are in the cerebral cortex, the more top-down predictions are inhibited. In other words, while in stress, neuromodulation systems amplifies the precision of sensory prediction errors, endowing sensory information with greater weight, in relation to prior expectations. As a consequence of chronic stress, individuals with high trait anxiety are unable to optimally update how volatile an aversive environment is, whereas low-anxiety controls exhibit close to Bayes optimum behavior (18).

Acute mental or psychosocial stress increases whole-brain glucose uptake by more than 10% (19). In essence, stress implies inflated uncertainty which, in turn, may lead to heightened rates of PE formation. The increment in gain of neurons encoding prediction errors is metabolically costly. When animals or people experience high uncertainty or feel threatened as a result of an unsteady nature of internal or external environment, they need to enter a hypervigilant state to downregulate the uncertainty as quick as possible (14). In order to gather the necessary information, additional cerebral energy is required. As will be discussed hereinafter, the mechanisms by which brain can raise glucose uptake under stress may be altered during brain-body interactions (20, 21).

It is worth noting that it is not the brain per se that adapts to environmental demands but the organism as a whole. Remarkably, multitude of studies has so far determined the crucial role of brain-body interactions in cognitive, emotional and behavioral functioning. In the rest of the paper, we will argue that computational psychiatry could be complemented by (and thus benefit from) inclusion of bodily systems as inseparable and indispensable component of cognition.

### Dialog between brain and body: computational perspective and relevance for stress

Not so long ago a raw of interesting models under an umbrella term of embodied cognition have been put forth to combine ideas about reciprocal and dynamical nature of relationships between brain, body, and environment (22). Put it more formally, embodied cognition is a set of theories emphasizing the role of body constitution and experience of interactions of the organism with environment in the functioning of various cognitive processes. Interested readers might be referred to (23) for further scrutiny.

Another piece of evidence comes from several theories wherein bodily signals have been proposed to play a major role in the emergence of emotionality and subjectivity (24, 25). Beyond just being a source of different transmitters and hormones or simply performing their mechanistic functions in periphery, it has been suggested the gut and heart may be considered as ticking clocks that constantly send information up to the central nervous system. Monitoring of those signals by the brain thereby create a self-centered neural reference frame from which first-person perspective can develop (26).

For the sake of brevity, the rest of this article will be mainly devoted to brain-gut interactions and implications for furthering computational psychiatry and stress research fields.

In recent times fascinating evidence has been provided in favor of existence of phase amplitude coupling in organismic level (27). In the study authors found that phase of the lower frequency gastric pacemaker was able to modulate the amplitude of alpha waves in the anterior insula, region widely engaged in stress, emotion, interception and self-awareness (28).

More specifically, detailed information regarding meal quality and/or quantity, or even types of nutrients ingested could be transmitted to the brain via vagal afferents. Gastric slow wave activity changes markedly in response to neurotransmitters and hormones, hence it is likely that the insular cortex persistently monitors such changes (29). However, the individual studies cannot portray a full picture of interactions between brain and body.

More recently, computational psychosomatics was put forward as an attempt to remedy this issue (30).

Authors of the paper underline that computational treatments of psychosomatic disorders have so far been rare. Mirroring trends in active adoption of dynamic systems approach in neuroscience, the term “allostasis” begin to be actively used (31). Allostasis means a readiness for changes through active adaptation (as opposed to homeostasis, described from the standpoint of maintaining of functioning mode in range of optimal values of physiological constants). Degree of impairment of the ability to adapt through changes is reflected in concept of allostatic load (32).

Allostatic load refers to the consequences or repeated activations of allostasis mediators, leading to inefficient functioning mode of allostatic system. Elevated levels of stress hormones after a certain time period can result in accumulation of allostatic load and pathophysiological consequences. Allostatic load markers can be divided into 4 subgroups: cardiovascular (diastolic and systolic blood pressure, resting heart rate), metabolic (body mass index, waist to hip ratio, total cholesterol, hemoglobin), inflammation (C-reactive protein), and stress-related hormones (epinephrine, norepinephrine, cortisol, dehydroepiandrosterone) (33).

Moving toward more integrative framework, predictive coding has been extended to account for allostatic perspective of brain-body interactions, which gave birth to embodied predictive interoception coding (EPIC) model, which unifies an anatomical model of corticocortical connections with Bayesian active inference principles (34). For example, from the standpoint of computational psychosomatics, false high-level beliefs about volatility of the world could cause prolonged allostatic responses (30).

However, different primary disturbances induce distinct patterns of change. Computational psychiatry and psychosomatics could catch this by statistical comparison of models entailing alternative disease processes. Combining predetermined perturbations with model selection and impending assessments of disease trajectories portray an encouraging approach to resolve ambiguity created by circular causality. However, by now there are only a few techniques for intentional probing the interoceptive circuits. These include cardiac challenges with short-acting sympathomimetics, manipulating inspiratory breathing load or air composition, transcutaneous vagus nerve stimulation, baroreceptor stimulation, acute induction of inflammation by vaccination, or C-fiber stimulation under capsaicin (30). We propose that this list might be complemented by adding techniques for dietary depletion of neuromodulators. The acute tryptophan depletion (ATD) and acute tyrosine plus phenylalanine depletion (ATPD) tests are powerful tools for studying contribution of cerebral monoamines to behavior, symptoms related to various disorders and improvement of treatment strategies (35, 36). For example, beverages lacking of tyrosine and phenylalanine may provide a unique way towards nutritional therapy of manic and psychotic disorders by inhibition of cerebral dopamine synthesis and release (37). Also, this could conceivably be used as an adjuvant treatment in patients who are resistant to conventional antipsychotic medication.

Moreover, predictive coding theory postulates that there exist, at least, two mechanisms to deal with priors when they need to be updated. First one is perceptual inference whereby the brain can update its internal model of the world by changing empirical prior beliefs (38). Another way to cope with imposed challenge is by leveraging active inference which aims at changing sensations by means of action or sequence of actions in order to make them more like predictions (39).

Active inference set a stage for understanding how our actions give rise to specific patterns of behavior, particularly under stress conditions. During stress, to cope with uncertainty we need to choose between approach and avoidance course of action (40).

Exploratory behavior is costly because it involves a variety of risks, leading to short-term uncertainty. Nevertheless, the potential merit of exploratory behavior is the lasting decline of uncertainty. In contrast, avoidance behavior can result in only transient uncertainty diminution and promotes adherence to outdated beliefs, thereby engendering long-term uncertainty (14). Several studies have conducted to find out an extent to which gastrointestinal system can influence animal’s preferred type of action. The results convincingly demonstrate that microbiota not only does have this influence but also that it may represent universal, conserved across species mechanism (41, 42).

The links between avoidance behavior and enterotypes of human gut microbiota are complicated and remain to be tested. However, it provides a unique opportunity for looking into the intricacies by targeting microbiota with probiotics, prebiotics or antibiotics under stress with concurrent noninvasive brain activity recordings. For example, EEG alpha rhythm asymmetry in frontal lobe serves as a well-known biomarker of approach-withdrawal tendencies and such an index can be used for longitudinal tracking dynamic nature of changes in approach-withdrawal behavior and its correlations with gut microbiota composition (43).

### Gut-microbiota-brain axis in regulation of brain computations

Recent advances have elucidated bidirectional nature of relationships between the enteric nervous system and brain. Gut-derived hormones have profound influence on brain activity. So far, two of the most investigated hormones implicated in mechanisms of hunger and satiety are ghrelin and leptin, respectively. In addition to maintenance of body weight, blood glucose and adiposity, ghrelin has been suggested to hold a key to several physiologic processes in the brain, such as neuroprotection, neurogenesis, anti-depressive and anti-anxiety mechanisms (44, 45). Leptin, in contrast, is gut peptide that regulate energy metabolism by exerting anorexigenic effect. Leptin inhibits the rewarding effects of food via mesolimbic reward circuits as well as hypothalamus (46). Previous results have shown that specific deletion of leptin receptors in glutamatergic neurons located in the hippocampus and prefrontal cortex causes a depressive-like phenotype, whereas ablation of the leptin receptors in dopamine neurons results in a robust anxiogenic phenotype (47).

One more example is neuropeptide Y (NPY), being a gut peptide, is widely expressed in the CNS and regulates physiological and behavioral responses such as stress and anxiety, fear, learning and memory, control of blood pressure, and sympathetic activity (48). NPY has gained attention as a stress resiliency transmitter associated with posttraumatic stress disorder (49).

Another key factor in maintenance of brain-gut cooperation is a gut microbiota which is a collection of different types of microorganisms inhabiting the human intestine. In recent years gut microbiota gain popularity and has been extensively studied as a risk factor for development of psychiatric and somatic disorders (50). The growing interest in intestinal inhabitants is reflected in dynamics of a number of publications in the PubMed database: at the request of “gut microbiota” one can find only 103 articles dated 2007, 700 articles were published in 2012 and up to 4110 articles in 2017. Main pathways through which microbiota may contribute to developing psychiatric disorders may be summarized as follows: modification of gut permeability, synthesis of neuropeptides and neurotransmitters, modulation of local and peripheral inflammation, reduced absorption of good nutrients, increased synthesis of potentially neurotoxic compounds, reduction of antioxidant status and increased lipid peroxidation, increased carbohydrate malabsorption, BDNF modulation (51).

It is of additional importance that intestinal microorganisms can independently synthesize a bunch of key neurotransmitters such as serotonin (Candida, Streptococcus, Enterococcus), GABA (Lactobacillus, Bifidobacteria, L. rhamnosus), norepinephrine (Escherichia, Bacillus), dopamine (Bacillus), acetylcholine (Lactobacillus) (51).

Gut microbiota alterations have been demonstrated across different mental and somatic disorders. High rates of gastrointestinal problems have been found in schizophrenia, partly as a result of alterations in immune system. Levels of Clostridiales, Prevotella and Lactobacillus ruminis in gut microbiota and level of choline in ACC are elevated in ultra-high risk subjects (52).

Recent studies provide complementary evidence that urbanization/ city living may causally contribute to development of schizophrenia and anxiety disorders (53, 54). It remains to be tested whether these lifestyle hallmarks may be linked to western type diet. This diet is notoriously associated with high-fat and high-sugar foods consumption (55).

Moreover, it has been shown that microbiota, acting through immune system, assures intestinal protection in mouse model of experimental stroke (56). Along the same line, another study argue that gut dysbiosis impairs recovery after spinal cord injury (57).

According to the present literature, restoration and enrichment of the intestinal microbiota affects reduction of depressive symptoms and anxiety, improves cognitive function and immunity. Indeed, higher intake of monounsaturated fats seems to be protective in patients with cognitive decline (58). Similarly, it has been shown that probiotics may represent a valuable aid in PTSD and chronic fatigue syndrome therapy (59, 60).

Modern science begins to uncover the crucial role of gut microbiota in development of Alzheimer’s and Parkinson’s diseases. To date, there already exist a few timely and thoughtful reviews and meta-analyses published on this issue (61, 62).

Therefore, targeting gut microbiota is promising strategy for treating wide range of disorders. Comparing microbiota alterations across disorders permit elicitation of common routes that may interplay with diverse environmental and genetic risk factors to generate disease-specific patterns of symptoms.

Studies also show that gut microbiota mediate relationship between reinforcing values of foods and midbrain dopamine (63). Rewarding properties of sweet palatable foods, in turn, can confer stress relief (64). However, overconsumption of sugar has been hypothesized to increase the risk of depression via several plausible biological pathways, where midbrain dopamine plays a key role (65, 66). Additional studies suggest that even brief exposure to obesogenic diet disrupts brain dopamine networks (67).

### Limitations and future directions

In this brief review, we strove to highlight some facets of the computational psychiatry approach which mainly engage in characterizing and measuring different kinds of brain’s inferences.

Mental health care system now makes active attempts to adapt and incorporate a raw of approaches developed under computational framework into clinical practice. The most vivid examples are computational psychiatry (including developmental computational psychiatry), computational psychosomatics, computational approach in psychotherapy (68), computational neurology (69) and neuropsychology (70), to mention only a few. This paper, in line with aforementioned directions of research, tries to take advantage of computational perspective to account for several previously unresolved questions.

For the most part, the article is focused on consideration of stress through lens of gut-microbiota-brain axis and possible ways for connecting it to computational psychiatry framework. This line of research should be complemented by investigating other body systems such as cardiovascular, respiratory, immune, reproductive system, with respect to their connections with central nervous system. Explicit testing of complex hypotheses and quantitative description of relations to be tested are among major strengths of the burgeoning research area.

We propose several prospective directions for future studies. Firstly, since gut microbiota composition can affect brain myelination patterns throughout animal and human development, impacting velocity of brain signals propagation (71). Noninvasive techniques such as EEG and stimulation methods like transcranial magnetic stimulation and transcutaneous vagus stimulation can represent valuable aid for investigating gut-brain interactions and devising testable hypotheses about relationships between axon myelination patterns and brain oscillations.

Analogously, since the vagus nerve represents a major route for brain-body communication, it may be of value to restore homeostasis or reduce allostatic load after stress by selectively targeting this pathway (72). By now, there are several studies arguing in favor of beneficial effects of transcutaneous vagus stimulation, leading to improvements in symptoms of depression and epilepsy (73, 74).

However, there are only a few studies that use EEG method in studying the phenomenon of microbiota by targeting it by means of pre-, pro-, and antibiotics. For example, A.P. Allen and his colleagues point to statistically significant improvements in hippocampus-dependent visuospatial memory performance after taking probiotics and enhanced frontal midline electroencephalographic mobility (75). Also, it has been reported that single treatment session with deep TMS over the PFC and insula improves symptoms of obesity by altering gut microbiota composition (76). One might suggest that gut microbiota enterotypes can be used concurrently with qEEG parameters to better inform prediction of treatment response by using machine learning techniques.

Likewise, one can argue that levels of beta oscillations provide a measure of the likelihood that a voluntary action will need to be actuated (77). Crucially, gut microbiota composition, type of diet or gut hormones (f.e. leptin) may be able to modulate beta activity (most likely, through dopamine-mediated mechanism), which reflects preparation of potential actions. Loss of dopamine, as in Parkinson’s disease, annuls this function. Thus, electrophysiological biomarkers can provide a useful tool for tracking brain-body interactions.

Secondly, we need to take into account developmental perspective of brain-body interactions, i.e. influence of early life adversities and sensitive periods.

For example, neglecting these sensitive developmental periods for the emergence of psychiatric disorders risks can hide the mechanisms that cause psychiatric disorders (78). A main goal of developmental computational psychiatry is to determine how developmental trajectories across multiple systems emerge and interact, and how deviations from the normative tracks eventually cause full-fledged disorder to appear.

Unfortunately, longitudinal developmental studies of computational tasks, used to probe computational brain mechanisms, are sparse. And there are almost no studies have investigated how psychiatric disorders are related to developmental aspects of these deficits (79).

Finally, another possible line of future research is to characterize extent to which gut microbiota composition and diet type can influence drug metabolism. This issue has far reaching implications for developing novel treatment strategies for psychiatric disorders. It has been reported that gut microbiota neuromodulators can cross blood-brain barrier (80) and microbiota, in general, can modulate the blood-brain barrier permeability through different routes (81). These findings may open a new way for manipulating brain-blood barrier properties in order to selectively and timely deliver specific drug into the brain.

Besides, traditional therapy of mental disorders with antidepressants, anxiolytics, neuroleptics, anticonvulsants does not always bring the desired or expected effect. A problem of side effects during long periods of drug intake is another significant difficulty for certain categories of population such as children, pregnant women, the elderly and patients with severe somatic diseases. Therefore, the development of new strategies for treatment of mental disorders will allow overcoming at least some of these hurdles. Ability of specific strains of gut microbiota to alleviate depressive, anxiety or cognitive symptoms has been repeatedly demonstrated over the past few years (56, 59). Microbiota targeting with pre- and probiotics or changes in diet can be also considered as an adjuvant for traditional therapy regimens.

Adopting developmental computational psychiatry and psychosomatics perspective to understanding intricate relationships between comaturation of HPA axis and gut microbiota will allow realizing the role played by their partnership during early life and adolescence sensitive periods. As we now know, these are timeframes during which animals or humans are highly susceptible to stressors’ influence (82).

Although promising, extant studies suffer from some limitations. For example, some surveys found no effects of probiotic treatment on symptoms of psychiatric disorders (83) and others argue that in healthy humans microbiota interventions with probiotics does not result in any changes of gut microbiota composition (84).

Taken together, results of ongoing studies is limited in their translational value because of small sample sizes, neglect of heterogeneity of psychiatric disorders, lack of randomization, insufficient duration of exposure.

Turning to advantages of proposed research directions, one might claim that from the computational psychiatry standpoint we could track specific trajectories in emergence of psychiatric symptoms starting with procedures of acute stress induction in lab settings in healthy controls, gradually moving towards full-blown major depression and anxiety disorders through a stage of subclinical manifestations of depression or anxiety. The subclinical level of disorders is one of the least studied fields in biological psychiatry with wide-ranging implications for predictions of treatment responses or searching for patients being at high risk for developing specific disorder.

In conclusion, computational psychiatry is rapidly developing research field, which provides important clues for elaboration of evidence-based personalized interventions, that is particular patients could be offered specific treatment regimens depending on points at which they diverge from theoretically optimal computational mechanisms. Modern big datasets hold a great promise to unmask these complex relations. We hope that this review will serve as an initial stepping-stone for constructing more integrated approaches to deal with psychiatric disorders in near future.

## Acknowledgments

The article was supported by research grant for the implementation of basic part of state assignment according to the Project 𝒩<u>o</u> 19.8259.2017/BCh.

## References

1. McEwen BS, Bowles NP, Gray JD, Hill MN, Hunter RG, Karatsoreos IN, Nasca C (2015): Mechanisms of stress in the brain. Nature neuroscience 18 (10): 1353–63.

2. Sapolsky RM (2015): Stress and the brain: individual variability and the inverted-U. Nature Neuroscience. 18 (10): 1344–6.

3. Koolhaas JM, Bartolomucci A, Buwalda BB, Boer SF, Flügge G, Korte SM., et al. (2011): Stress revisited: a critical evaluation of the stress concept. Neuroscience and biobehavioral reviews 35 (5): 1291–301.

4. Montague PR, Dolan RJ, Friston KJ, Dayan P (2012): Computational psychiatry. Trends in cognitive sciences 16 (1): 72–80.

5. Huys QJ, Maia TV, Frank MJ (2016): Computational psychiatry as a bridge from neuroscience to clinical applications. Nature Neuroscience 19: 404–413.

6. Friston KJ, Redish AD, Gordon JA (2017): Computational Nosology and Precision Psychiatry. Computational Psychiatry 1: 2–23.

7. Weinberger DR, Glick ID, Klein DF (2015): Whither Research Domain Criteria (RDoC)? The Good, the Bad, and the Ugly. JAMA psychiatry 72 (12): 1161–2.

8. Kirmayer LJ, Crafa DA (2014): What kind of science for psychiatry? Front. Hum. Neurosci. 8: 435.

9. McGorry PD, Keshavan MS, Goldstone SD, Amminger PG, Allott KA, Berk M, et al. (2014): Biomarkers and clinical staging in psychiatry. World psychiatry: official journal of the World Psychiatric Association 13 (3): 211–23.

10. Dwyer DB, Falkai P, Koutsouleris N (2018): Machine Learning Approaches for Clinical Psychology and Psychiatry. Annual review of clinical psychology (14): 91–118.

11. Bzdok D, Meyer-Lindenberg A (2018): Machine Learning for Precision Psychiatry: Opportunities and Challenges. Biological psychiatry. Cognitive neuroscience and neuroimaging 3 (3): 223–230.

12. Bastos AM, Usrey WM, Adams RA, Mangun GR, Fries P, Friston KJ (2012): Canonical Microcircuits for Predictive Coding. Neuron 76: 695–711.

13. Bubic A, von Cramon DY, Schubotz RI (2010): Prediction, cognition and the brain. Frontiers in human neuroscience 4: 25.

14. Peters A, McEwen BS, Friston K (2017): Uncertainty and stress: Why it causes diseases and how it is mastered by the brain. Progress in Neurobiology 156: 164–188.

15. de Berker AO, Rutledge RB, Mathys C, Marshall L, Cross GF, Dolan RJ, Bestmann S (2016): Computations of uncertainty mediate acute stress responses in humans. Nature communications 7: 10996.

16. Ulrich-Lai YM, Herman JP (2009): Neural regulation of endocrine and autonomic stress responses. Nature Reviews Neuroscience 10: 397–409.

17. Yu AJ, Dayan P (2005): Uncertainty, Neuromodulation, and Attention. Neuron 46: 681–692.

18. Browning M, Behrens TE, Jocham G, O’Reilly JX, Bishop SJ (2015): Anxious individuals have difficulty learning the causal statistics of aversive environments. Nature neuroscience 18(4): 590–6.

19. Hitze B, Hubold C, Dyken RV, Schlichting K, Lehnert H, Entringer S, Peters A (2010): How the Selfish Brain Organizes its Supply and Demand. Front. Neuroenerg. 2: 7

20. Mergenthaler P, Lindauer U, Dienel GA, Meisel A (2013): Sugar for the brain: the role of glucose in physiological and pathological brain function. Trends in neurosciences, 36(10): 587–97.

21. Lee SH, Zabolotny JM, Huang H, Lee H, Kim YB (2016): Insulin in the nervous system and the mind: Functions in metabolism, memory, and mood. Molecular metabolism 5(8): 589–601.

22. Lynott D, Connell L, Holler J (2013): The role of body and environment in cognition. Front. Psychol. 4: 465.

23. Pfeifer R, Iida F, Lungarella M (2014): Cognition from the bottom up: on biological inspiration, body morphology, and soft materials. Trends in cognitive sciences 18 (8): 404–13.

24. Craig AD (2010): The sentient self. Brain Structure and Function 214: 563–577.

25. Dunn BD, Dalgleish T, Lawrence AD (2006): The somatic marker hypothesis: a critical evaluation. Neuroscience and biobehavioral reviews 30 (2): 239–71.

26. Park H, Tallon-Baudry C (2014): The neural subjective frame: from bodily signals to perceptual consciousness. Philosophical transactions of the Royal Society of London. Series B, Biological sciences 369 (1641): 20130208.

27. Richter CG, Babo-Rebelo M, Schwartz D, Tallon-Baudry C (2017): Phase-amplitude coupling at the organism level: The amplitude of spontaneous alpha rhythm fluctuations varies with the phase of the infra-slow gastric basal rhythm. NeuroImage 146: 951–958.

28. Craig AD (2009): How do you feel — now? The anterior insula and human awareness. Nature Reviews Neuroscience 10: 59–70.

29. Huizinga JD (2017): Commentary: Phase-amplitude coupling at the organism level: The amplitude of spontaneous alpha rhythm fluctuations varies with the phase of the infra-slow gastric basal rhythm. Frontiers in neuroscience 11: 102.

30. Petzschner F, Weber LA, Gard T, Stephan KE (2017): Computational Psychosomatics and Computational Psychiatry: Toward a Joint Framework for Differential Diagnosis. Biological Psychiatry 82: 421–430.

31. Sterling P (2012): Allostasis: a model of predictive regulation. Physiology & behavior 106: 5–15.

32. Karatsoreos IN, McEwen BS (2011): Psychobiological allostasis: resistance, resilience and vulnerability. Trends in cognitive sciences 15 (12): 576–84.

33. Savransky A, Chiappelli J, Fisseha F, Wisner KM, Xiaoming D, Mirmomen SM, et al. (2018): Elevated allostatic load early in the course of schizophrenia. Translational psychiatry 8(1): 246.

34. Barrett LF, Simmons WK (2015): Interoceptive predictions in the brain. Nature reviews. Neuroscience 16(7): 419–29.

35. Faulkner P, Deakin JF (2014): The role of serotonin in reward, punishment and behavioural inhibition in humans: insights from studies with acute tryptophan depletion. Neuroscience and biobehavioral reviews 46: 365–78.

36. Ramdani C, Vidal F, Dagher A, Carbonnell L, Hasbroucq T (2018): Dopamine and response selection: an Acute Phenylalanine/Tyrosine Depletion study. Psychopharmacology 235: 1307–1316.

37. Badawy AA 2013) Novel nutritional treatment for manic and psychotic disorders: a review of tryptophan and tyrosine depletion studies and the potential of protein-based formulations using glycomacropeptide. Psychopharmacology 228: 347–358.

38. Aggelopoulos NC (2015): Perceptual inference. Neuroscience and biobehavioral reviews 55: 375–92.

39. Friston KJ, FitzGerald TH, Rigoli F, Schwartenbeck P, O’Doherty JP, Pezzulo G (2016): Active inference and learning. Neuroscience & Biobehavioral Reviews 68: 862–879.

40. Stein MB, Paulus MP (2009): Imbalance of approach and avoidance: the yin and yang of anxiety disorders. Biological psychiatry 66 (12): 1072–4.

41. Lee K, Mylonakis E (2017): An Intestine-Derived Neuropeptide Controls Avoidance Behavior in Caenorhabditis elegans. Cell reports 20 (10): 2501–2512.

42. Szyszkowicz JK, Wong A, Anisman H, Merali Z, Audet M (2017): Implications of the gut microbiota in vulnerability to the social avoidance effects of chronic social defeat in male mice. Brain, behavior, and immunity 66: 45–55.

43. Adolph D, Glischinski MV, Wannemueller A, Margraf J (2017): The influence of frontal alpha-asymmetry on the processing of approach- and withdrawal-related stimuli-A multichannel psychophysiology study. Psychophysiology 54 (9): 1295–1310.

44. Andrews ZB (2011): The extra-hypothalamic actions of ghrelin on neuronal function. Trends in neurosciences 34 (1): 31–40.

45. Wittekind DA, Kluge M (2015): Ghrelin in psychiatric disorders - A review. Psychoneuroendocrinology 52: 176–94.

46. Fulton S, Woodside B, Shizgal P (2000): Modulation of brain reward circuitry by leptin. Science 287 (5450): 125–8.

47. Liu J, Guo M, Lu XY (2015): Leptin/LepRb in the Ventral Tegmental Area Mediates Anxiety-Related Behaviors. The international journal of neuropsychopharmacology, 19 (2): 1–11.

48. Eaton K, Sallee FR, Sah RA (2007): Relevance of neuropeptide Y (NPY) in psychiatry. Current topics in medicinal chemistry 7 (17): 1645–59.

49. Sah RA, Geracioti TD (2013): Neuropeptide Y and posttraumatic stress disorder. Molecular Psychiatry, 18, 646–655.

50. Cryan JF, Dinan TG (2012): Mind-altering microorganisms: the impact of the gut microbiota on brain and behavior. Nature Reviews Neuroscience 13: 701–712.

51. Sherwin E, Dinan TG, Cryan JF (2018). Recent developments in understanding the role of the gut microbiota in brain health and disease. Annals of the New York Academy of Sciences: 1420 (1): 5–25.

52. He Y, Kosciólek T, Tang J, Zhou Y, Li Z, Ma X, et al. (2018): Gut microbiome and magnetic resonance spectroscopy study of subjects at ultra-high risk for psychosis may support the membrane hypothesis. European psychiatry: the journal of the Association of European Psychiatrists 53: 37–45.

53. Lederbogen F, Kirsch P, Haddad L, Streit F, Tost H, Schuch P, et al. (2011): City living and urban upbringing affect neural social stress processing in humans. Nature (474): 498–501.

54. Lambert KG, Nelson RJ, Jovanovic T, Cerda ME (2015): Brains in the city: Neurobiological effects of urbanization. Neuroscience and biobehavioral reviews 58: 107–22.

55. Jacka FN (2017): Nutritional Psychiatry: Where to Next? EBioMedicine 17: 24–29.

56. Winek K, Dirnagl U, Meisel A (2016): The Gut Microbiome as Therapeutic Target in Central Nervous System Diseases: Implications for Stroke. Neurotherapeutics 13 (4): 762–774

57. Kigerl KA, Hall JC, Wang L, Mo X, Yu Z, Popovich PG (2016): Gut dysbiosis impairs recovery after spinal cord injury. The Journal of experimental medicine 213 (12): 2603.

58. Morris MC, Tangney CC, Wang Y, Sacks FM, Barnes LL, Bennett DA, Aggarwal NT (2015): MIND diet slows cognitive decline with aging. Alzheimer’s & dementia: the journal of the Alzheimer’s Association 11 (9): 1015–22.

59. Gilbert JA, Krajmalnik-Brown R, Porazinska DL, Weiss S, Knight R (2013): Toward Effective Probiotics for Autism and Other Neurodevelopmental Disorders. Cell 155: 1446–1448.

60. Campagnolo N, Johnston S, Collatz A, Staines D, Marshall-Gradisnik S (2017): Dietary and nutrition interventions for the therapeutic treatment of chronic fatigue syndrome/myalgic encephalomyelitis: a systematic review. Journal of human nutrition and dietetics: the official journal of the British Dietetic Association 30 (3): 247–259.

61. Borre YE, O’Keeffe GW, Clarke G, Stanton C, Dinan TG, Cryan JF (2014): Microbiota and neurodevelopmental windows: implications for brain disorders. Trends in molecular medicine 20 (9): 509–18.

62. Sampson TR, Debelius JW, Thron T, Janssen S, Shastri GG, Ilhan ZE, et al. (2016): Gut Microbiota Regulate Motor Deficits and Neuroinflammation in a Model of Parkinson’s Disease. Cell 167: 1469–1480.

63. Volkow ND, Wise R, Baler RD (2017): The dopamine motive system: implications for drug and food addiction. Nature Reviews Neuroscience 18: 741–752.

64. Ulrich-Lai YM, Fulton S, Wilson ME, Petrovich GD, Rinaman L (2015): Stress exposure, food intake and emotional state. Stress 18 (4): 381–99.

65. Knüppel A, Shipley MJ, Llewellyn CH, Brunner EJ (2017): Sugar intake from sweet food and beverages, common mental disorder and depression: prospective findings from the Whitehall II study. Scientific reports 7 (1): 6287.

66. Hu D, Cheng L, Jiang W (2019) Sugar-sweetened beverages consumption and the risk of depression: A meta-analysis of observational studies. Journal of Affective Disorders 245: 348–355.

67. Barry RL, Byun NE, Williams JM, Siuta MA, Tantawy MN, Speed NK, et al. (2018): Brief exposure to obesogenic diet disrupts brain dopamine networks. PloS One 13 (4): e0191299.

68. Moutoussis M, Shahar N, Hauser TU, Dolan RJ (2018): Computation in Psychotherapy, or How Computational Psychiatry Can Aid Learning-Based Psychological Therapies. Computational Psychiatry 2: 50–73.

69. Freund P, Friston K, Thompson AJ, Stephan KE, Ashburner J, Bach DR, et al. (2016): Embodied neurology: an integrative framework for neurological disorders. Brain: a journal of neurology 139 (6): 1855–61.

70. Parr T, Rees G, Friston KJ (2018): Computational Neuropsychology and Bayesian Inference. Front. Hum. Neurosci. 12: 61.

71. Hoban AE, Stilling RM, Ryan FJ, Shanahan F, Dinan TG, Claesson MJ, et al. (2016). Regulation of prefrontal cortex myelination by the microbiota. Translational Psychiatry. 6 (4): e774.

72. Mayer EA (2011): Gut feelings: the emerging biology of gut–brain communication. Nature Reviews Neuroscience 12: 453–466.

73. Fang J, Egorova N, Rong P, Liu J, Hong Y, Fan Y, et al. (2017). Early cortical biomarkers of longitudinal transcutaneous vagus nerve stimulation treatment success in depression. NeuroImage: Clinical 14: 105–111.

74. Helmers SL, Duh MS, Guérin A, Sarda SP, Samuelson TM, Bunker MT, et al. (2011). Clinical and economic impact of vagus nerve stimulation therapy in patients with drug-resistant epilepsy. Epilepsy & behavior: E&B 22 (2): 370–5.

75. Allen AP, Hutch W, Borre YE, Kennedy P, Temko A, Boylan G, et al. (2016). Bifidobacterium longum 1714 as a translational psychobiotic: modulation of stress, electrophysiology and neurocognition in healthy volunteers. Translational psychiatry. 6 (11): e939.

76. Ferrulli A, Macrì C, Terruzzi I, Ambrogi F, Milani V, Adamo M et al. (2018): High frequency deep transcranial magnetic stimulation acutely increases β-endorphins in obese humans. Endocrine: In press.

77. Jenkinson N, Brown P (2011): New insights into the relationship between dopamine, beta oscillations and motor function. Trends in neurosciences 34 (12): 611–8.

78. Frankenhuis WE, Nettle D, McNamara JM (2018): Echoes of Early Life: Recent Insights From Mathematical Modeling. Child development 89 (5): 1504–1518.

79. Hauser TU, Will GJ, Dubois M, Dolan RJ (2018): Developmental Computational Psychiatry. J Child Psychology & Psychiatry

80. Bonaz B, Bazin T, Pellissier S (2018): The Vagus Nerve at the Interface of the Microbiota-Gut-Brain Axis. Frontiers in neuroscience 12: 49.

81. Kelly JR, Minuto C, Cryan JF, Clarke G, Dinan TG (2017): Cross Talk: The Microbiota and Neurodevelopmental Disorders. Front. Neurosci. 11: 490.

82. Oitzl MS, Champagne DL, Veen RV, Kloet ER (2010): Brain development under stress: hypotheses of glucocorticoid actions revisited. Neuroscience and biobehavioral reviews 34 (6): 853–66.

83. Reis DJ, Ilardi SS, Punt SE (2018): The anxiolytic effect of probiotics: A systematic review and meta-analysis of the clinical and preclinical literature. PloS One. 13 (6): e0199041.

84. Kristensen NB, Bryrup T, Allin KH, Nielsen T, Hansen TH, Pedersen O (2016): Alterations in fecal microbiota composition by probiotic supplementation in healthy adults: a systematic review of randomized controlled trials. Genome Medicine. 8 (1): 52.

